# Towards understanding the regulation of histone H1 somatic subtypes with OMICs

**DOI:** 10.1101/2020.09.30.320572

**Authors:** Inma Ponte, Marta Andrés, Albert Jordan, Alicia Roque

**Affiliations:** Biochemistry and Molecular Biology Department, Bioscience Faculty, Autonomous University of Barcelona, Spain; Molecular Biology Institute of Barcelona (IBMB-CSIC), Barcelona, Spain

**Keywords:** co-regulation, transcription factors, compensatory effects, subtype functional differentiation, chromatin compartments, topologically associated domains (TADs)

## Abstract

**Background:** Histone H1 is involved in the regulation of chromatin higher-order structure and compaction. In humans, histone H1 is a multigene family with seven subtypes differentially expressed in somatic cells. Which are the regulatory mechanisms that determine the variability of the H1 complement is a long-standing biological question regarding histone H1. We have used a new approach based on the integration of OMICs data to address this question.

**Results:** We have examined the 3D-chromatin structure, the binding of transcription factors (TFs), and the expression of somatic H1 genes in human cell lines, using data from public repositories, such as ENCODE. Analysis of Hi-C, ChIP-seq, and RNA-seq data, have shown that transcriptional control has a greater impact on H1 regulation than previously thought. Somatic H1 genes located in TADs show higher expression than in boundaries. H1 genes are targeted by a variable number of transcription factors including cell cycle-related TFs, and tissue-specific TFs, suggesting a finetuned, subtype-specific transcriptional control. We describe, for the first time, that all H1 somatic subtypes are under transcriptional co-regulation. The replication-independent subtypes, which are encoded in different chromosomes, isolated from other histone genes are also co-regulated with the rest of the somatic H1 genes, indicating that transcriptional co-regulation extends beyond the histone cluster.

**Conclusions:** Transcriptional control and transcriptional co-regulation explain, at least in part, the variability of H1 complement, the fluctuations of H1 subtypes during development, and also the compensatory effects observed, in model systems, after perturbation of one or more H1 subtypes.

## Background

Histone H1 is involved in the regulation of chromatin structure and transcriptional regulation. In humans, histone H1 is a multigene family encoding eleven subtypes or variants, which are grouped into seven somatic subtypes, and four germ-line-specific subtypes [1]. The latter group includes one female germ-line subtype and three male germ-line-specific subtypes. Somatic cells express variable amounts of the remaining seven subtypes, H1.0-H1.5 and H1x. The complement of histone H1, defined as the subtypes and its proportions present on a specific cell, is variable. It depends on the cell type, cell cycle phase and the time of development. However, the regulation of H1 complement in physiological conditions and disease is not fully understood.

Subtypes H1.1-H1.5 are encoded in chromosome 6, within the major histone cluster, which is found in the nucleus in a membrane-less, self-organizing compartment called histone locus body (HLB) [2]. Genes encoding histone H1 subtypes are single-copy genes, while there are multiple copies of core histone genes. HLB contains histone mRNA biosynthetic factors and its assembly has been suggested to be mediated by phase transition [3]. During S-phase, phosphorylation by CDK2 of NPAT coordinates the increase in the expression of histone genes, including those somatic H1 genes in the chromosome 6 cluster. Therefore, subtypes H1.1-H1.5 are known as replication-dependent subtypes (RD-subtypes). Transcripts of RD-subtypes are not polyadenylated. They have a conserved stem-loop structure in the 3’ UTR, which is involved in their post-transcriptional regulation [2]. The other two somatic subtypes, H1.0 and H1x, are encoded in chromosomes 22 and 3, respectively. Their transcription is uncoupled from HLB transcription and thus, they are known as replication-independent subtypes (RI-subtypes). Their transcripts are longer than those of the RD-subtypes and are also polyadenylated [4].

At a functional level, H1 somatic subtypes exhibit some redundancy as shown by knockout and knockdown experiments. Mice, where one or two somatic subtypes were inactivated, showed normal development and fertility [5, 6, 7, 8]. The ratio H1:core histones in these knockouts was normal, due to an increase in the remaining subtypes, mainly in H1.0. However, mouse embryos lacking three H1 subtypes (H1.2, H1.3, and H1.4) died by mid-gestation with a broad range of defects [8], while embryonic stem cells derived from this triple knockout showed impaired differentiation [9]. In triple knockouts the ratio H1:core histones was significantly reduced. It seems that subtype redundancy is directly associated with the basic structural function of H1, the maintenance of chromatin higher-order structure.

Compelling evidence of subtype functional differentiation has been found by knockdown experiments. Rapid inhibition of each H1 variant by sh-RNA, in human cell lines, is not compensated by changes in expression of other H1 variants. The reduction of H1.2 or H1.4 caused defects in proliferation in breast cancer cells [10], while the knockdown of H1.0 in human ESCs did not affect self-renewal but impaired differentiation [11]. H1.5 depletion in human fibroblasts caused decreased cell growth, increased chromatin accessibility, and alterations in gene expression [12]. Combined depletion of H1.2 and H1.4, in breast cancer cells, resulted in strong interferon response and induced a compensatory effect in H1, characterized by an increase in H1.0 expression [13].

Additional evidence also suggests that H1 subtypes are functionally distinct. Some examples are the subtype-specific genomic localization [12, 14, 15, 16], different chromatin binding-affinity [17, 18, 19], post-translational modifications [20], and interaction partners of individual subtypes [21]. Molecular evolution studies have also shown that H1 subtypes are under strong negative selection and that they display different evolution velocities [22, 23]. Additionally, it has been shown that positive selection events may have been involved in subtype diversification and in determining the low chromatin binding affinity of H1.1 [24].

The development and widespread use of OMIC techniques based on next-generation sequencing methods have generated large amounts of genomic/transcriptomic data that remains unexplored at the gene level. We have taken advantage of publicly available data to explore the 3D-chromatin organization of somatic H1 genes in human cell lines finding conserved features and also differential features. Also, we have shown that the location of H1 genes in TADs or boundaries is associated with their expression level. Analysis of ChIP-seq data has shown that the proximal promoters of somatic H1 genes were targeted by a wide variety of transcription factors (TFs) including general TFs, ligand-activated TFs, differentiation-associated TFs, and cell cycle-related TFs, suggesting a tight and subtype-specific transcriptional control. This analysis also revealed overlap in the TFs mapped to somatic H1 gene promoters, suggesting transcriptional coregulation. This hypothesis was supported by the correlation analysis of RNA-seq data, which found positive and negative correlations among individual subtypes. Additionally, we have also shown that only data from total RNA is reliable to analyze H1 expression levels.

## Results

### 3D-chromatin organization of the somatic H1 genes in human cell lines

The development of the Hi-C technique has allowed the quantification of the interaction between chromatin fragments at a genome-wide scale. Analysis of Hi-C data has shown the spatial segregation of open and closed chromatin to form two compartments, A and B, corresponding to active chromatin and inactive chromatin, respectively [25]. Analysis of chromatin compartments of 10 human cell lines from ENCODE has shown that genes encoding somatic H1s, located in chromosomes 3, 6 and 22, were always found in the A compartment (Figure 1, Figure 2A, Supplementary Table 1). The length of the A compartments was variable between chromosomes and cell lines (Supplementary Figure 1, Supplementary Table 2). However, the A compartments, where genes encoding H1.0 and H1x are located, were mostly conserved among the cell lines analyzed, as they had similar length and showed a high degree of overlapping in the pairwise intersection analysis (Figure 2B and 2C). Genes encoding subtypes H1.1-H1.5, located in the histone cluster in chromosome 6, were mapped within the same A compartment (Figure 1, Figure 2D). In this case, the length of this compartment was variable, with lower Jaccard coefficients than the compartments where H1.0 and H1x are located, indicating less degree of overlapping among the different cell lines (Figure 2D).

**Figure 1.**
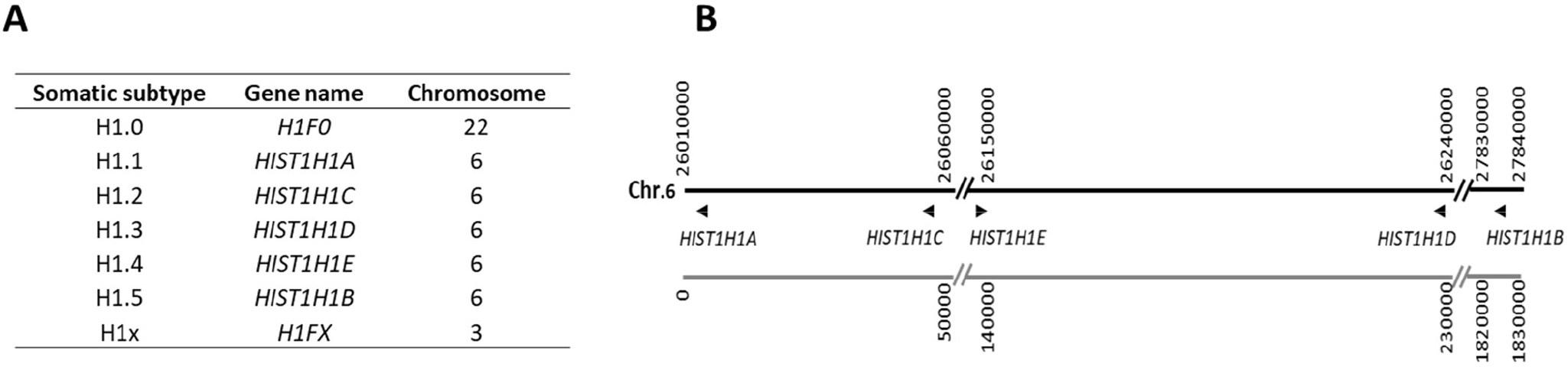
H1 somatic subtypes in humans. A, correspondence between H1 subtypes, gene name and genomic location. For alternative nomenclatures used for H1 somatic subtypes refer to Talbert et al. [1]. B, graphical representation of the H1 somatic subtypes genes located at the histone cluster in chromosome 6. The arrowheads indicate the orientation of each gene. The start and end positions of each chromosomal segment are shown. The diagonal double bars separate different segments of the chromosome. The positions of the genes are referred to *hg19* human genome assembly. The relative positions of the chromatin fragments are represented in the grey line.

**Figure 2.**
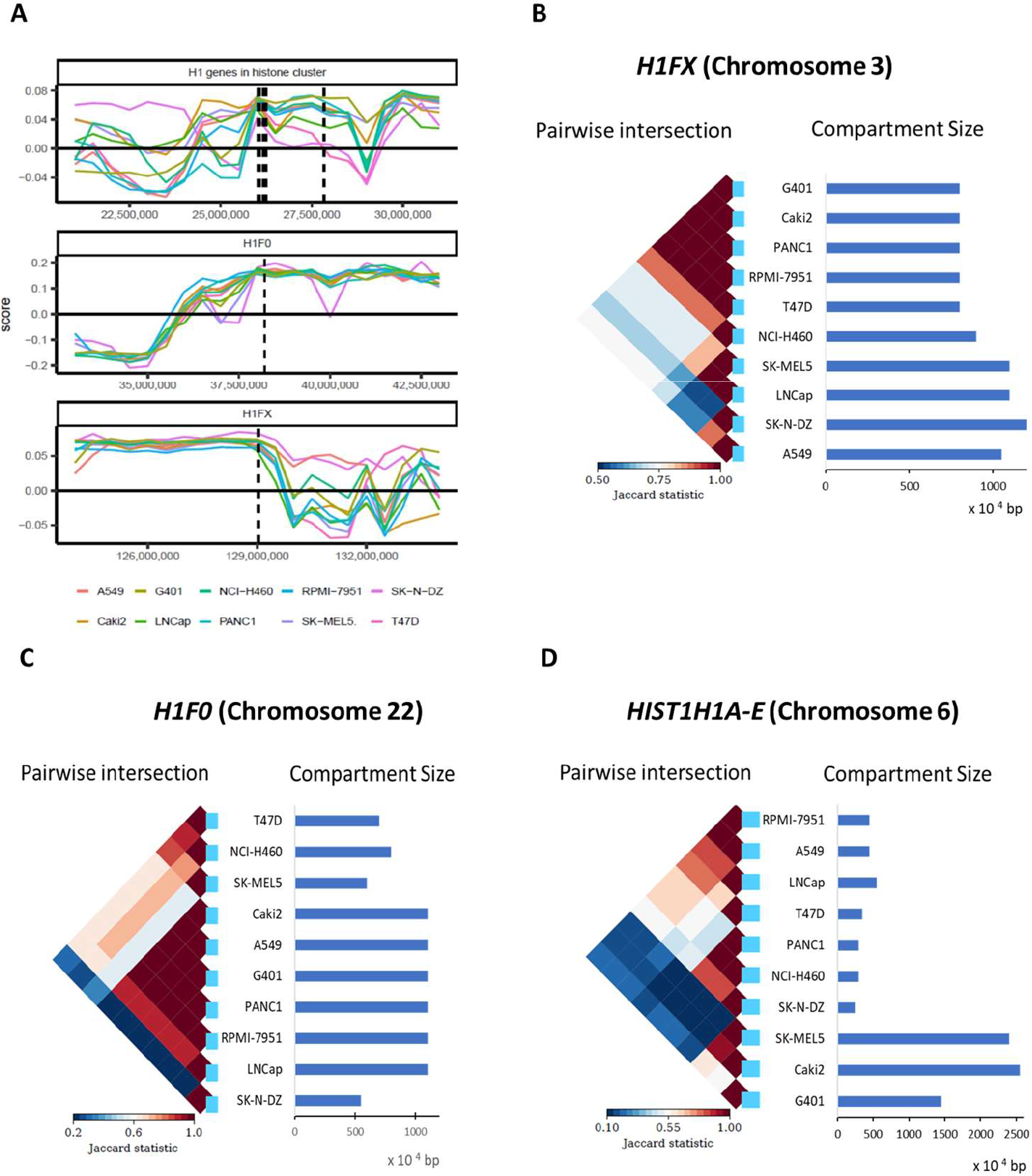
Compartments containing somatic H1 genes. A, Compartment analysis of the genomic region spanning 10 Mb containing H1 genes in chromosome 6 (top panel), in chromosome 22 (middle panel) and in chromosome 3 (lower panel). The lines correspond to the specific location of the H1 genes within the compartment. Pairwise intersection showing the overlapping in chromosomes 3 (B), 22 (C) and 6 (D) between the genomic region assigned to the A compartment containing H1 genes in different cell lines. The blue bars represent the length of the compartment in each cell line. The positions of the genes are referred to *hg19* human genome assembly.

In GM12878, compartments A and B have been further divided into 6 subcompartments (A1, A2, B1, B2, B3, and B4) based on their Hi-C interaction patterns [26]. These subcompartments also exhibit different genomic and epigenomic content. Genes encoding somatic H1 subtypes were mapped to the A1 subcompartment. This subcompartment is characterized by early replication, high gene content, the presence of highly expressed genes, and the enrichment in activating chromatin marks such as H3K36me2 and H3K27ac, among others.

Analysis of Hi-C contact matrices has also revealed that the genome is partitioned in regions of ~1 Mb called, topologically associated domains (TADs) [27]. The increase in the resolution of Hi-C experiments has resulted in the description of smaller contact regions called subTADs and even smaller contact domains of an average length of 185 kb [26], giving rise to a more general definition of TADs, accounting for contact regions between 100 kb and 1 Mb, approximately [28].

We have explored the 3D-organization of the chromatin regions, where the genes encoding H1 somatic subtypes are located. Information of 22 Hi-C experiments, which describe the chromatin 3D-structure of 21 different cell lines had been used [26, 27, 28, 29]. In three of the four datasets, chromatin was digested with *Hind* III, which recognition site spans six nucleotides [27, 28, 29]. Rao and colleagues used an *in situ* Hi-C protocol and *Mbo* I as restriction endonuclease, which is a 4-cutter enzyme to analyze eight different cell lines, IMR90, K562, GM12878, HMEC, HUVEC, HeLa, NHEK, and KBM7 [26]. The latter procedure has two main advantages, the reduction of the frequency of spurious contacts and the detection of smaller contact domains, than those observed with a 6-cutter enzyme. In this study, the highest resolution was obtained for the cell line GM12878, due to higher sequencing depth. In the four datasets, the TAD caller used allowed for inter-TADs gaps, generally termed boundaries, with variable or fixed length (Supplementary Table 3). Region of chromosomes 3, 6, and 22, encoding somatic H1 genes were analyzed in detail (Figure 3). The different resolution of the four Hi-C datasets resulted in the significant differences in the number of TADs annotated per megabase, as well as in the length of the TADs, in the three chromosomes encoding somatic H1 genes (Supplementary Figure 2).

**Figure 3.**
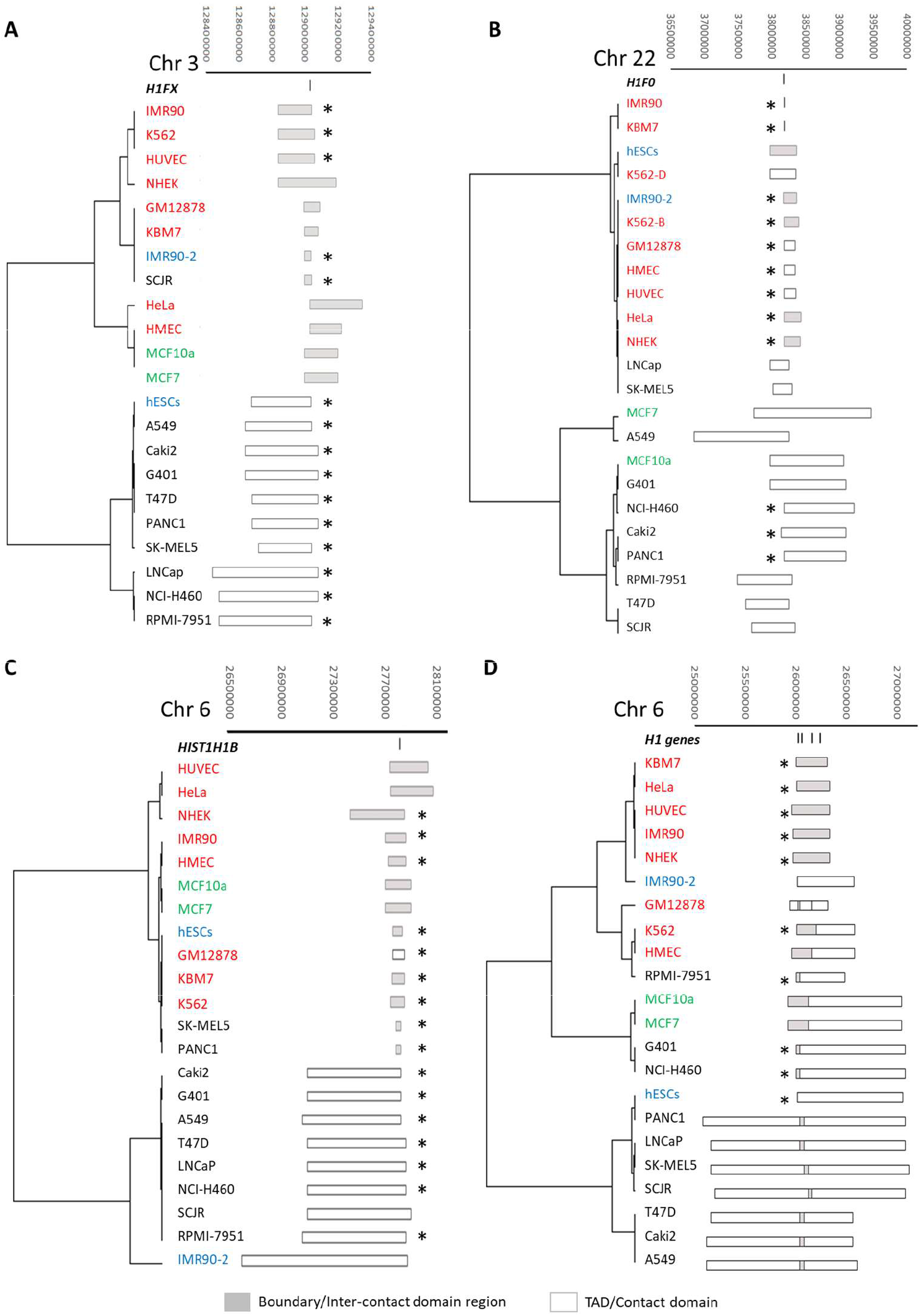
Chromatin 3D-organization of somatic H1 genes in human cell lines. A, *H1FX* gene. B, *H1F0* gene. *C, HIST1H1B* gene. D, *HIST1H1A, HIST1H1C-E* genes. Cell lines were ordered by hierarchical clustering of the limits of the chromatin regions. The upper panel shows the coordinates of the chromatin regions in the corresponding chromosome fragment. Lines in the first row denote the exact position of somatic H1 genes. Boxes represent chromatin regions where the H1 genes are located. Filled boxes correspond to boundaries or inter-contact domain regions, while empty boxes correspond to TADs or contact domains. All the positions are referred to *hg19* human genome assembly. Cell lines are colored according to the dataset of origin: in blue, dataset 1 [27]; in red, dataset 2 [26]; in black, dataset 3 [29] and in green, dataset 4 [28] Conserved borders are marked with asterisks (*).

Gene *H1FX*, encoding H1x, is located in the q-arm of chromosome 3. Results of Hi-C experiments show that *H1FX* gene can be either located within TADs or in boundary regions (Figure 3A, Supplementary Table 4). Despite the heterogeneity in the definition of the boundaries in the four datasets, they were shorter than TADs with an average size of 180 kb and 456 kb, respectively (Figure 4A). It was also apparent that both borders of the domains were not conserved, with only two genes, *HMCES* and *H1FX-AS*, always mapped together with *HIFX*. However, in 16 cell lines, from three different datasets, the 3’ border was conserved, even when the *H1FX* gene was located in a boundary or a TAD (Figure 3A). In all cell lines, except NHEK, the gene encoding H1x was less than 50 kb from the border of the domain.

**Figure 4.**
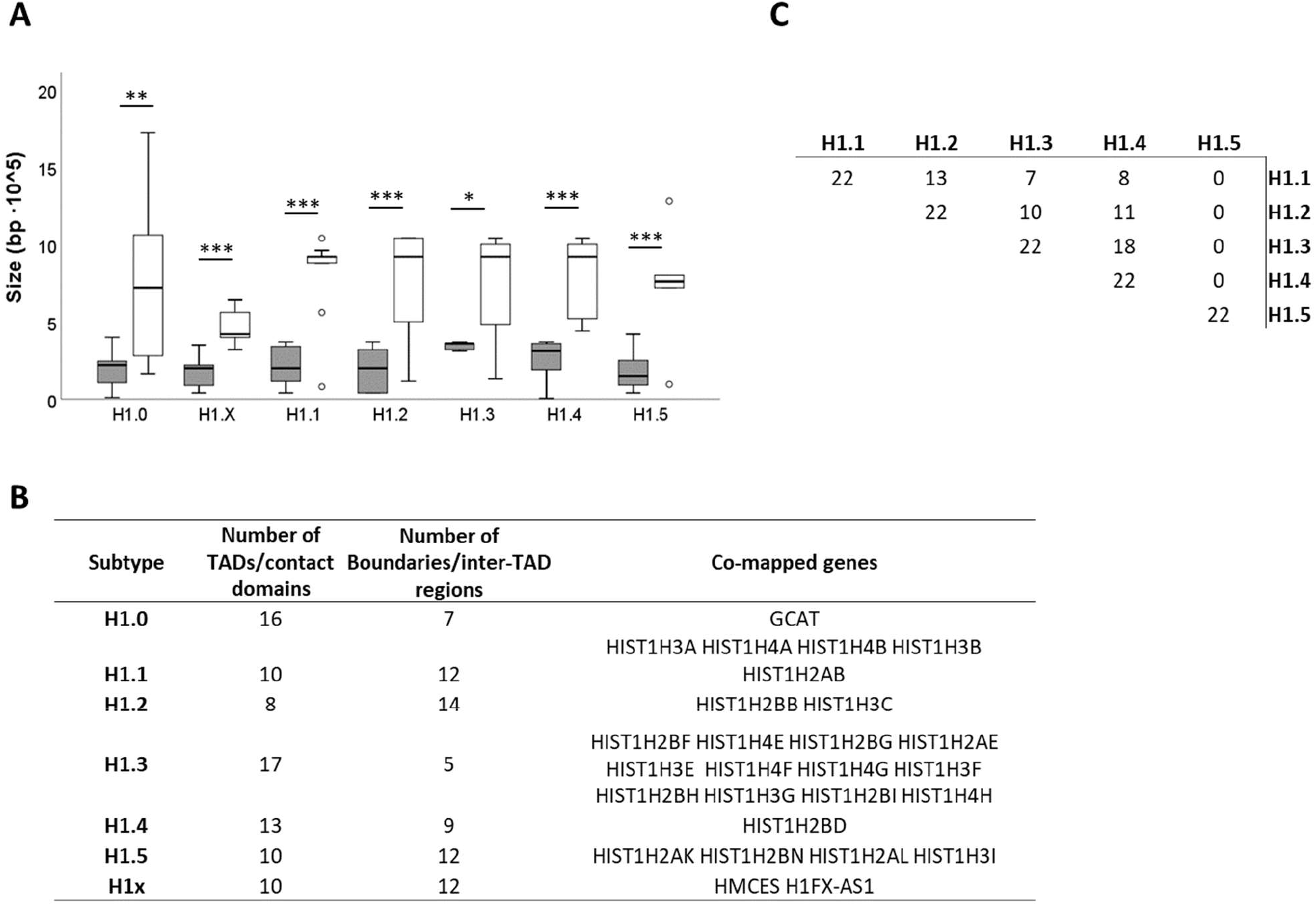
Analysis of the chromatin 3D-organization of somatic H1 genes in human cell lines. A, Box-plot of the size of regions containing each H1 somatic subtype, separated by type. Gray boxes correspond to boundaries, while empty boxes correspond to TADs. Asterisks correspond to the p-values obtained after comparing the size of boundaries and TADs using Mann-Whitney nonparametric test: *, p-value < 0.05; **, p-value < 0.01; ***, p-value < 0.001. B, Summary of the region type for individual H1 genes and of lines between the H1 somatic genes located at the histone cluster. The values in the matrix correspond to the number of domains or boundaries regions shared by a pair of H1 genes.

Gene *H1F0*, encoding H1.0, is located in the q-arm of chromosome 22. This gene has been mapped within TADs in 16 cell lines and in boundaries in seven cell lines, with an average size of 736 kb and 190 kb, respectively (Figure 3B, Figure 4A, Supplementary Table 5). The chromatin 3D-structure of this region, similar to that of the *H1FX* gene was variable, although the 5’ border was conserved in 12 cell lines, including cell lines from three different experimental studies. In those cell lines, the *H1F0* gene was less than 50 kb from the border of the domain. In two cell lines, IMR90 and KBM7, the H1.0 gene was mapped in a very small boundary of 10 kb. In this region, there were only two genes present *H1F0* and *GCAT*, which encodes an enzyme involved in L-threonine degradation. Therefore, *GCAT* was the only gene always mapped with *H1F0* (Figure 3B). In K562, a cell line derived from a chronic myeloid leukemia patient (CML), the *H1F0* gene was mapped in two different chromatin domains, in the same Hi-C experiment, one boundary and one TAD, indicating that the chromosomal alterations typical of this disease triggered changes in the 3D-chromatin organization of specific genes.

Genes encoding the replication-dependent subtypes H1.1-H1.5 are located in the histone cluster in the short-arm of chromosome 6, interspersed with core histone genes. Hi-C experiments have shown that H1.5, which is located more distal to the rest of the H1 genes present in the cluster, was mapped to a different chromatin region in all cell lines (Figure 1B, Figure 3C, Supplementary Table 6). This gene was mapped within TADs, in ten cell lines with an average size of 682 kb, and in boundaries, in 12 cell lines with a size of 166 kb on average (Figure 4A, Figure 4B). Despite the type of region, the 3’ border was conserved in 16 cell lines, in which the gene encoding H1.5 was less than 50 kb from the border of the domain (Figure 3C). The shortest domain, a boundary of 40 kb, containing the gene encoding H1.5 was found in melanoma and pancreatic cancer cell lines, SK-MEL5 and PANC1, respectively. In this region, four genes encoding core histones were found.

The rest of the H1 genes in the histone cluster were mapped within the same chromatin region in seven cell lines, two of them classified as TADs, and the other five as boundaries (Figure 3D, Supplementary Table 6). In GM12878, the cell line with the highest resolution, the four genes were mapped in different regions. In the rest of the cell lines, the genes encoding H1.1-H1.4 were partitioned in different domains, with genes encoding H1.3 and H1.4 being the two genes sharing the same chromatin domain in most cell lines (Figure 3D). In general, the genes encoding H1.1 and H1.2 were mapped more often in boundaries, while the genes encoding H1.3 and H1.4 were mapped more frequently in TADs (Figure 4B). The 5’ border of the region containing the gene encoding H1.1 was conserved in ten cell lines, with fluctuations of less than 90 kb in another five cell lines (Figure 3D). Only a few genes were always mapped with a specific H1 gene, due to the variability of the annotated chromatin regions (Figure 4B).

We have analyzed data from IMR90 derived from two different experiments [26, 27]. *H1F0* and *H1FX* were mapped in the same region type, a boundary region, that had one end conserved, while the other end differed, resulting in different sizes of the chromatin regions (Figure 3A, 3B). In the case of the H1 genes belonging to the histone cluster, the length and type of region were different between the experiments, while one of the ends of the defined chromatin regions was conserved (Figure 3C and 3D). The changes observed in the same cell line can be attributed to the different resolutions in the Hi-C experiments. In fact, significant differences were observed between the two datasets in the number of TADs/Mb and TADs length of the three chromosomes analyzed (Supplementary Figure 2).

### Binding of transcription factors to somatic H1 gene promoters

Early studies have described the topology of histone H1 promoters [30]. Two sequence-specific regions located within 1kb of the transcription start site (TSS), called H1-box and TG-box, have been shown to be important for H1 gene expression. These two elements are widely conserved in sequence and position, and they could be identified in the proximal promoters of all human somatic H1 genes, except in the gene encoding H1.1 (Supplementary Figure 3).

Chromatin immunoprecipitation coupled to next-generation-sequencing (ChIP-seq) has allowed obtaining genome-wide maps of chromatin-bound proteins. ChIP-seq analysis has also shown the genomic distribution of histone post-translational modifications, including those frequently found in active or silent gene promoters. Active promoters are enriched in H3K4me3 and H3K27ac, among others. In human cell lines, somatic H1 promoters were enriched in H3K27ac, especially those subtypes with highest mRNA levels H1.2 and H1.4, while low or moderate enrichment in H3K4me3 was detected (Supplementary Figure 4). Promoters of the RI-subtypes showed less abundance of H3K27ac than promoters of the RD-subtypes. Treatment of breast cancer cells, T47D with Trichostatin A, a histone deacetylase inhibitor (HDACi), resulted in the increase of the transcript levels of both, H1.0 and H1x, indicating that core histone acetylation can regulate their expression (Supplementary Figure 5). In contrast, the transcript levels of the RD-subtypes expressed in T47D showed very little changes. These results were confirmed in Jurkat cells (data not shown).

We have used the Gene Transcription Regulation Database (GTRD), which provides a collection of uniformly analyzed ChIP-seq data to get information about transcription factors binding close to the TSS of somatic H1 genes [31]. This database includes information regarding many chromatin-bound proteins, including remodeling complexes, chromatin-modifying enzymes, and transcription factors (TF). For further analysis, we have considered only those proteins listed as human transcription factors [32].

Between 64 and 279 TFs binding within 1kb upstream or downstream the TSS of somatic H1 genes were found (Figure 5A, Supplementary Table 7). The gene with less TFs mapped was the one encoding H1.1, while the gene with more TFs was the one encoding H1.4. Considerable overlapping was found in the four subtypes with more TFs bound to their promoters, which shared more than 130 TFs (Figure 5B). These four subtypes included, the two RD-subtypes expressed in most cell types, H1.2 and H1.4, and the two RI-subtypes, H1.0 and H1x.

**Figure 5.**
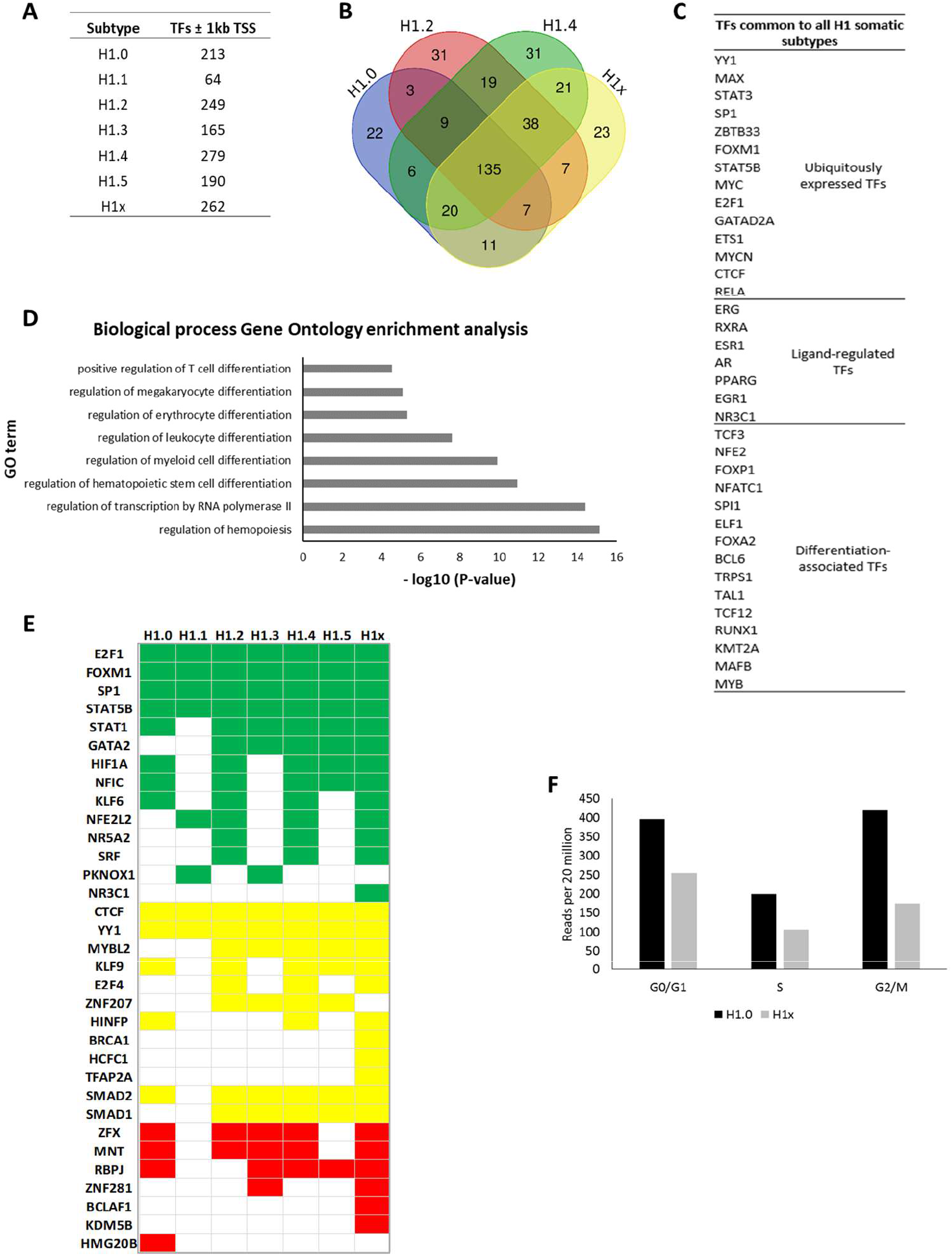
Transcriptional control of H1 gene promoters. A, Number of transcription factors mapped to H1 somatic gene promoters by ChIP-seq. B, Venn diagram of the 4 subtypes with more TFs detected in their gene promoters. C, Transcription factors common to all H1 somatic subtypes. D, Biological process Gene Ontology enrichment analysis of the differentiation-associated TFs. The enrichment analysis was performed using Panther GO enrichment online tool. For comparison, the GO term with lower p-value associated with transcription was included in the figure. For hematopoiesis, only GO terms with FDR <0.001 were considered. E, binding to individual somatic H1 gene promoters of cell-cycle-related TFs, and of previously described transcription factors affecting histone expression. In green, transcriptional activators; in red, transcriptional repressors, and in yellow, factors capable of activating and inhibiting transcription. F, levels of nascent RNA of the genes encoding H1.0 and H1x, obtained by GRO, during cell cycle in MCF7 (GSE94479).

At least 36 TFs were mapped to the promoters of all somatic H1 genes, which could be classified as ubiquitously expressed TFs, ligand-activated TFs and differentiation-associated transcription factors (Figure 5C). Interestingly, the last category was enriched in TFs associated with the differentiation of myeloid and lymphoid cell-types (Figure 5D). In fact, 11 out of the 15 differentiation-associated TFs are directly involved in hematopoiesis regulation.

A detailed analysis of TFs associated with cell cycle [33], as well as TFs previously reported bound to histone genes [2, 34, 35] was performed. We found that many cell-cycle-associated TFs have been mapped to H1 gene promoters, including known transcriptional activators, transcriptional repressors, and transcription factors, which may either activate or repress their target genes, depending on other conditions (Figure 5E). Some of these TFs, including SP1 and E2F1, were mapped to all somatic H1 promoters. SP1 has been proposed to bind the GC-rich element present in H1 promoters [30]. E2F1 has been associated with the activation of G1/S gene expression [36] and has also been identified as a master regulator of histone expression in mouse ESCs [34].

In addition to E2F1, several of the main transcription factors controlling the periodical gene expression during the cell cycle have been mapped to H1 gene promoters. This TF group includes E2F4, MYBL2 (b-myb) and FOXM1 [35] (Figure 5E). E2F4 was mapped to the proximal promoters of three of the seven somatic subtypes (H1.2, H1.4, and H1x), while MYBL2 was mapped to five H1 gene promoters and FOXM1 appeared to bind all H1 gene promoters. The binding of cell cycle transcriptional regulators to H1 gene promoters is in agreement with data form genomic run on (GRO) experiments in MCF7 synchronized cells, which have confirmed *de novo* transcription of the RI-subtypes, during all phases of the cell cycle, including G2/M [37] (Figure 5F). Data regarding *de novo* transcription of H1 RD-subtypes was not available, because only polyadenylated transcripts were analyzed.

Furthermore, transcription factors, previously described as having a role in histone transcription, were also mapped in several H1 human genes. H1NFP has been mapped to 3 H1 genes, H1.0, H1.4, and H1x (Figure 5E). This TF can activate or repress its target genes. H1NFP has been described to activate transcription of H4, and to repress the transcription of retinoblastoma protein 1, RB1 [35, 38]. YY1 and CTCF were mapped to all H1 gene promoters. CTCF has been shown to negatively correlate with histone expression in mouse ESCs, while the binding of YY1 to histone promoters has been proposed to be restricted to mouse differentiated cells [34]. SMAD1, SMAD2, and ZNF have been mapped to five, six, and five H1 gene promoters, respectively (Figure 5E). These TFs have been described as negative regulators of histone transcription in mouse ESCs [34].

It has been shown that HBP1, a partner of the retinoblastoma protein (RB) activates *H1F0* transcription in murine erythroleukemia cells (MEL), linking this subtype to cell cycle [39]. This TF was mapped to H1.0, as well as H1.2 and H1.4 proximal promoters in human cell lines (Supplementary Table 7). H1.0 has also been described to respond to external signals, in particular in glands requiring hormonal stimulation in mice (reviewed in [40]. This characteristic is apparently conserved in human cell lines, where several hormone receptors were mapped to H1.0 proximal promoter. Knockout of FOXA2 in mouse embryos reduced the transcript levels of H1x [41]. In human cell lines, this TF has also been mapped to the proximal promoter of H1x, as well as to the rest of somatic H1 genes (Figure 5C). These results suggest that transcriptional regulation of H1 genes is conserved in mammalians.

### Transcriptomic data of H1 subtypes

One of the most prolific OMIC is transcriptomic, which has generated an incredible amount of data describing the transcriptional landscape of multiple cell types of different species in normal or in specific conditions. Great consortiums like Genotype Tissue Expression (GTEx) (https://www.gtexportal.org/) and Cancer Cell Line Encyclopedia (CCLE), among others, have been created as an effort to provide transcriptomic information from human tissues and cell lines, whose results are accessible at the projects website’s or have been compiled into public portals like the EBI’s Expression Atlas (https://www.ebi.ac.uk/gxa/home). In addition, transcriptomic data generated by independent scientific research is, for the most part, readily available at public repositories such as the Gene Expression Omnibus (GEO) (https://www.ncbi.nlm.nih.gov/gds).

At the experimental level, most of the transcriptomic data is derived from microarray and RNA-seq experiments, where transcripts of protein-expressing genes have been selected by affinity procedures based on their polyA tails. Somatic H1 transcripts are heterogenous, H1.0 and H1x have polyadenylated mRNAs, while the rest of the subtypes lack polyA tail. For that reason, in the case of somatic H1 genes, the majority of the transcriptomic data is not accurate.

As a proof of concept, we have analyzed the available RNA-seq data of H1 somatic subtypes in CCLE, which contains more than one thousand human cancer cell lines [42, 43]. Despite the heterogeneity of the data, the average expression values for each subtype showed that the more represented subtypes were those with polyadenylated mRNAs, H1.0 and H1x. H1.2 was the most abundant, among RD-subtypes, although the reason why this subtype is preferentially purified in polyA-selected RNA-seq experiments is unknown (Figure 6A).

**Figure 6.**
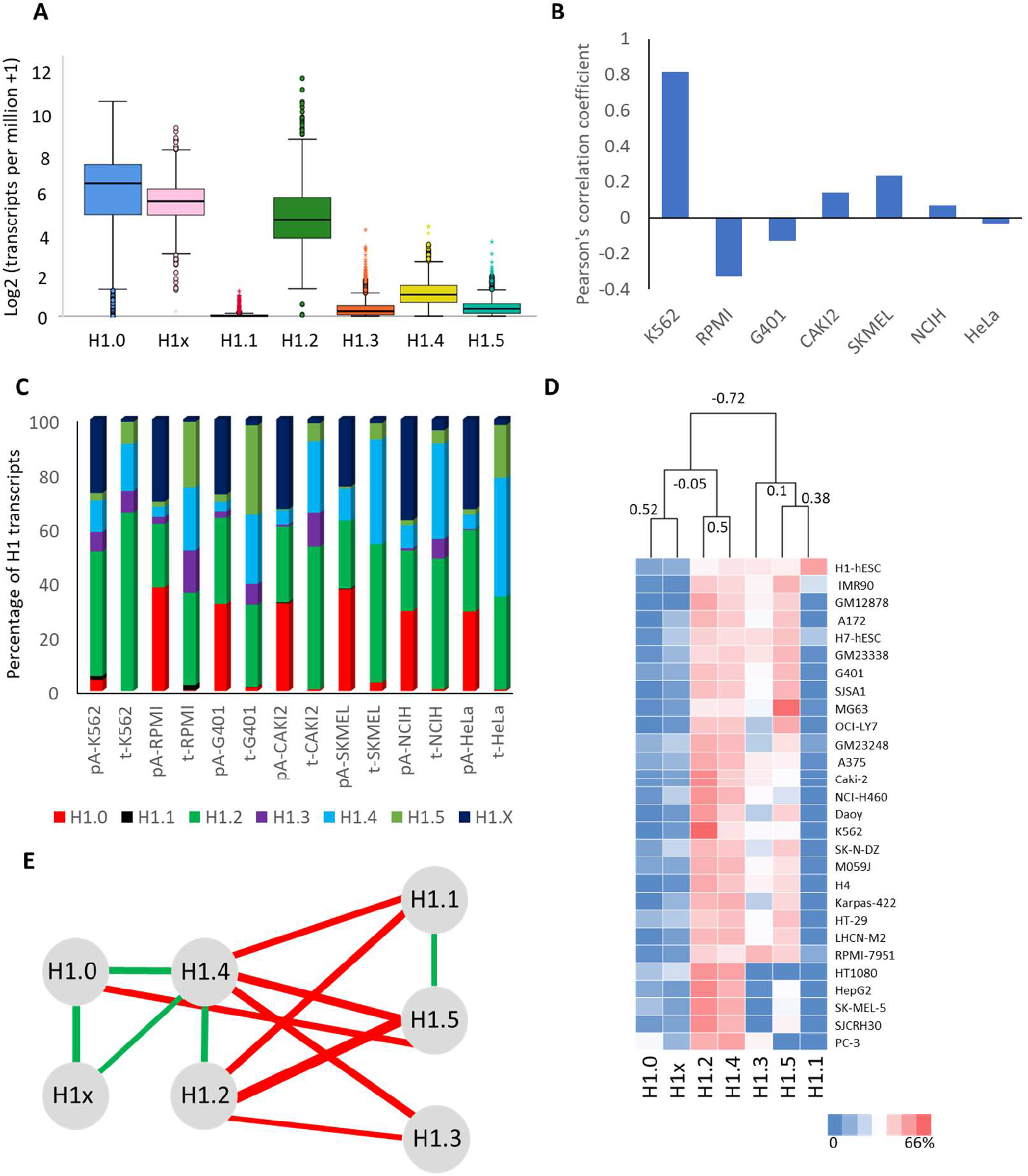
Transcriptomic data of H1 somatic subtypes. A, Box plot of the transcript levels of H1 somatic subtypes from the CCLE [42, 43]. B, Pearson’s correlation coefficients of somatic H1 transcript levels detected in RNA-seq selecting polyadenylated transcripts and total RNA-seq experiments, in human cell lines. C, Contribution in percentage of each subtype to the total H1 transcripts detected in RNA-seq from polyadenylated RNA (pA-RNA) or total RNA (t-RNA). D, Hierarchical clustering analysis based on the Spearman rank correlation coefficient. The numbers of the dendogram indicate the similarity distance for each node. E, Correlation network for the expression of H1 subtypes. In green, positive correlation coefficients; in red negative correlation coefficients. The width of each line corresponds to the absolute value of the correlation coefficient. Only correlation coefficients with p-values < 0.05 were represented. The data analyzed corresponded to all RNA-seq experiments (from total RNA) of human cell lines, available at ENCODE [29]. Abundance of each transcript was expressed as percentage of all the transcripts encoding H1 somatic subtypes.

Comparison of the mRNA relative abundance by subtype, between RNA-seq based on polyA selection and RNA-seq from total RNA, of a small subset of cell lines, resulted in low or even negative Pearson’s correlation coefficients between the two datasets (Figure 6B). However, high Pearson’s correlation coefficients could be obtained occasionally, depending on the specific composition of the H1 complement of the cell line. Drastic changes in the proportions of the subtype transcripts could be observed (Figure 6C). When polyadenylated RNA was purified, between 77-93% of the H1 coding transcripts corresponded to H1.0, H1.x, and H1.2. In contrast, when total RNA was analyzed the same subtypes represented between 33-66% of the total H1 transcripts. The more significant change was the contribution of subtypes H1.0 and H1x to the total number of H1 transcripts, whose combined proportions decreased from 30-60% when polyadenylated transcripts were selected, to less than 5% in RNA-seq data from total RNA. These results indicate that only transcriptomic data derived from total RNA is reliable for the analysis of H1 transcript levels.

Analysis of the transcriptome data may also facilitate the study of possible transcriptional coregulation in H1 somatic subtypes. For this purpose, we have used RNA-seq data derived from total RNA samples of 28 human cell lines. The hierarchical clustering of H1 somatic subtypes based on Spearman rank correlation resulted in the separation of the seven subtypes in two clusters (Figure 6D). The first cluster was formed by H1.0, H1.2, H1.4, and H1x. Within this cluster, subtypes were further separated into two subclusters, one containing the RD-subtypes H1.2, and H1.4, and the other containing the RI-subtypes. Subtypes H1.1, H1.3, and H1.5 were assigned to the second cluster. In this case, the distance between subtypes H1.1 and H1.5 was shorter than the distance with H1.3.

Based on the pairwise Spearman’s rank correlation coefficients, a correlation network was constructed (Figure 6E, Supplementary Table 8). The transcript levels among the four subtypes within the first cluster were positively correlated. Interestingly, this cluster is formed by the four subtypes, whose genes were targeted by more TFs, which showed ~ 50% of overlap. Thus, it appears that transcriptional co-regulation is responsible, at least in part, for the correlations at the mRNA level. In the second cluster, only the pair H1.1-H1.5 showed a positive correlation, while the transcript levels of all subtypes within this cluster were negatively correlated with the transcript levels of H1.2 and H1.4. Our results suggest that despite additional effects due to post-transcriptional processing, co-regulation at transcriptional level may play a role in the regulation of the H1 complement.

Finally, we wanted to know whether the 3D-chromatin structure of H1 genes was associated with their expression levels among human cell lines. We used transcriptomic data from total RNA of nine human cell lines, where the 3D-chromatin organization had been described. We found that when H1 genes were in TADs or contact domains their relative expression was higher that when they were in boundaries (Figure 7A). This tendency was observed in all H1 subtypes, except H1.1. The expression of H1.1 is restricted to certain tissues and its general expression is quite low. Other exceptions were observed in H1.0, H1.3, H1.4, and H1.5. In H1.4, the highest relative expression was found in GM12878, where the gene encoding this subtype was in a boundary, whereas in the other three subtypes the lowest expression was when the genes were in TADs (Figure 7A). Overall, in 84% of the cases, H1 genes with high relative expression in different human cell lines were located in TADs, whereas only 16% were located in boundaries (Figure 7B), suggesting a clear relationship between chromatin 3D-structure and the expression of H1 genes.

**Figure 7.**
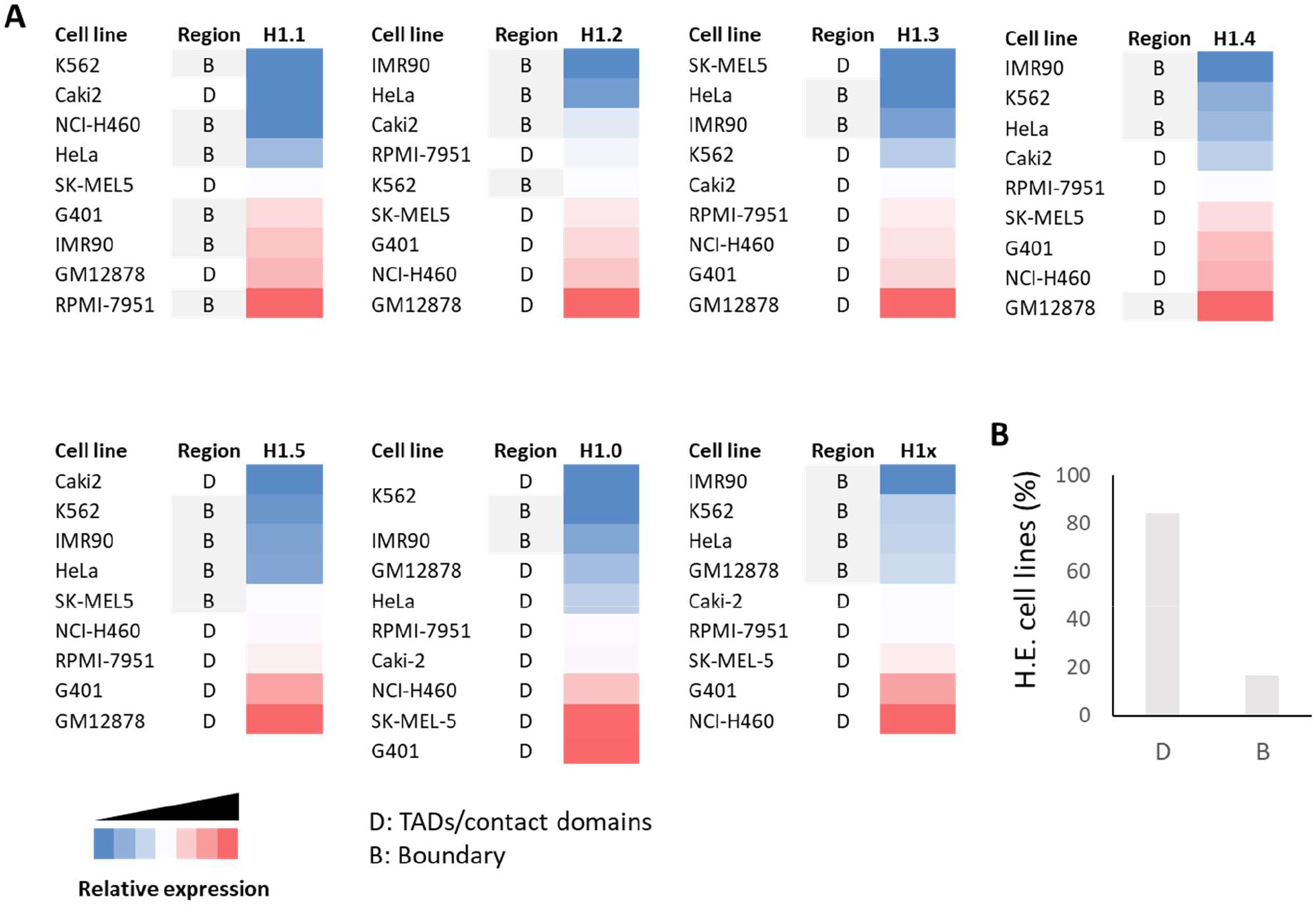
Association of 3D-chromatin structure with the expression level of H1 subtypes. A, Correspondence between the relative expression levels and type of chromatin region in human cell lines. For IMR90, the study with the highest resolution was selected. K562 was not considered in H1.0, due to the presence of the gene encoding this subtype in a TAD and a boundary, simultaneously. Relative expression values were calculated after normalization to GAPDH. B, Overall percentages of the type chromatin region found in cell lines with high expression (H.E. cell lines) by subtype.

## Discussion

Genes encoding somatic H1 subtypes are located in three different chromosomes. Replication dependent subtypes, H1.1-H1.5, are encoded in chromosome 6, within the major histone cluster, while replication-independent subtypes, H1.0 and H1x, are encoded in chromosomes 22 and 3, respectively. Analysis of Hi-C data showed that, in human cell lines, H1 genes were located in the active chromatin compartment (A compartment). In GM12878, where 6 subcompartments were defined, all somatic H1 genes were mapped in the A1 compartment, which replicates early in S-phase, contains highly expressed genes and it’s enriched in active chromatin marks like H3K27ac [26]. Chromatin regions with the genes encoding H1.0 and H1x, assigned to compartment A, were more conserved, among human cell lines, than the region of chromosome 6 containing genes encoding H1 RD-subtypes.

Close examination of the 3D-chromatin organization of the genomic regions encoding H1 somatic subtypes has shown that H1 genes have been mapped to chromatin regions of variable size, defined either as TADs or boundaries. It has been suggested that TADs and boundaries are different at a functional level. While chromatin regions within TADs interact with each other, regions located at TAD boundaries have been shown to be enriched in housekeeping genes, specific core histone modifications, and CTCF [27]. In addition, genes in boundaries are usually highly transcribed, suggesting that high levels of transcription activity may also contribute to boundary formation. It remains to be explored if the presence of H1 genes in different chromatin regions affects their expression.

Very few genes were mapped with individual somatic H1 genes in all cell lines. However, in all the regions analyzed, at least one of the borders of the chromatin domain was conserved in several cell lines from different Hi-C datasets. In addition, genes encoding subtypes H1.0, H1x, and H1.5 were close to the conserved borders in most of the cell lines. The differences observed in the 3Dchromatin structure of H1 genes can be attributed to several factors: i) experimental procedure and algorithms used for data processing, ii) changes typical of cancer cell lines, iii) heterogeneity of domain organization in a cell population, and iv) chromatin dynamics. It has to be noted, that in this comparison hierarchical TAD organization, which may result in nested or overlapping TADs was not considered.

Footprints of the reciprocal translocation between chromosomes 9 and 22, typically found in chronic myeloid leukemia [44], were detected by Hi-C analysis of the cell line K562, derived from a patient with this disease. In this cell line, the gene *H1F0*, which is located in the region of chromosome 22 affected by the translocation, was mapped in two different chromatin regions (one TAD and one boundary), indicating alterations in chromatin interactions caused by its presence on different chromosomes.

Previous studies of the 3D-chromatin organization of the HLB in breast cancer cell lines at different stages of cancer progression have shown that the histone cluster was divided into three TADs with the histone genes located at the boundaries [45]. This organization left the gene encoding H1.5 in the boundary of a TAD, separated from the rest of H1 coding genes. It was also shown that the boundaries interacted with each other more frequently than with intra-TAD regions, suggesting the formation of an active chromatin hub [45]. Analysis of this region in 21 human cell lines confirmed that the gene encoding H1.5 is located at a different chromatin region than the rest of H1 genes. However, the 3D-organization of the histone cluster in human cell lines was variable, as H1 encoding genes were found either at boundaries or TADs of variable length and gene composition. However, the interactions between the regions encoding histone genes may be preserved despite the variability of its 3D-chromatin organization by the formation of an active chromatin hub within the HLB. The variability of this region is also apparent at compartment level, and it may be related somehow to the formation of the histone locus body.

The 3D-chromatin organization has been implicated in transcriptional regulation [26, 46, 47]. It is thought that most of the promoter-enhancer interactions take place within individual TADs, and therefore TADs restrict the nuclear search space of regulatory elements [46, 47]. Our analysis showed that the 3D-chromatin organization of H1 genes is associated with their relative expression. We found that most H1 genes had a higher relative expression when they were located within TADs or contact domains, suggesting the presence of intra-TAD promoter-enhancer interactions.

It has been described that transcript levels of the genes encoding H1.1-H1.5, located at the histone cluster in chromosome 6, are increased during S-phase. Their up-regulation during replication is coupled to that of core histone genes and coordinated by phosphorylation of NPAT by CDK2 [2]. However, key TFs recognizing DNA elements within H2B and H4 promoters necessary for their transcription have been described [48, 49, 50]. In contrast, for somatic H1 genes, no key TF has been described, even if their transcript levels are variable in different cell types.

Global transcriptional control during the cell cycle is carried out basically by a few transcription factors: E2F factors, MYBL2, and FOXM1 [36]. Repressor E2Fs, including E2F4, are components of the DREAM complex in G0, which is involved in the repression of all cell cycle-dependent gene expression. Activators E2F factors, which include E2F1, are recruited to the promoters of genes whose expression peak during the G1 to S phase transition. Finally, MYBL2 and FOXM1 are sequentially recruited to cell cycle gene promoters by MuvB during S-phase and G2, to G2/M gene promoters.

ChIP-seq data showed that E2F4 has been mapped to some H1 genes, but not to the H1.0 promoter. This finding agrees with previous results describing that H1.0 transcription increased during cell arrest in G0 [10, 13, 51]. Unexpectedly, the knockout of E2F4 in mouse fibroblasts decreased the expression of several histone genes, including H1.2 [34]. E2F1 has been mapped to all somatic H1 genes, accordingly to its previously assigned role as a master regulator of H1 transcription during S-phase [34]. Surprisingly, TFs related to expression during G2/M, MYBL2 and FOXM1 have also been mapped to somatic H1 genes. In particular, FOXM1 is associated with strong gene activation during G2/M and was mapped in all somatic H1 genes [52]. Recent studies have shown that mitotic chromatin is far from inert at the transcriptional level as some transcription factors, as well as transcription machinery, remain bound to chromatin in mitosis [53]. As a result, thousands of genes are transcribed during mitosis, albeit at lower levels than in interphase, due to the lack of enhancer-promoter contacts [37]. *H1F0* and *H1FX* were among the genes, whose nascent transcripts were detected during G2/M in synchronized MCF7 cells by GRO [37]. It remains to be explored if the rest of somatic H1 genes are also transcribed at this cell cycle phase.

ChIP-seq analysis has also identified other TFs related to cell cycle bound to all somatic H1 promoters, suggesting combinatorial effects in their transcriptional control. Thus, all somatic H1 subtypes, including H1.0 and H1x, seem to be under tight and gene-specific transcriptional control throughout the cell cycle. Additionally, several transcription factors detected by ChIP-seq in human cell lines, including Smad1, YY1, HBP1, and FOXA2, had been already identified experimentally in mouse tissues or cell lines, suggesting that H1 regulation is conserved among mammalian species [34, 39, 41].

H1 complement is variable among different cell types and also during development. We have found that several TFs associated with the regulation of hematopoiesis were mapped to all H1 promoters, suggesting the importance of H1 for this process. Our finding is in agreement with recent data showing that epigenetic silencing of the gene encoding H1.3 is associated with a lower aggressive phenotype of acute myeloid leukemia (AML), suggesting that this subtype plays a role in AML blast differentiation [54]. Previous data showing that dendritic cells differentiation was significantly decreased in H1.0 knockout mice, also supports our conclusion [55]. In a very recent publication, several H1 subtypes have been associated with neutrophil differentiation [56]. Disruption of the expression of H1 subtypes with CRISPR-Cas 9 has shown that loss of H1.2 and H1.4 inhibited neutrophil differentiation and increased the expression of some eosinophil proteins. In contrast, deficiency in H1.1, H1.3, and H1.5 enhanced neutrophil differentiation. It would be interesting to uncover more evidence of the role of H1 in the differentiation of hematopoietic lineages or in leukemia.

Analysis of the transcription factors mapped to H1 proximal promoters by ChIP-seq has shown that the number of TFs targeting the TSS proximal region was associated with the capability of a somatic H1 gene to be expressed in different cell types or conditions. Accordingly, the gene with less TFs was the one encoding H1.1, which corresponds with the subtype with more restricted expression in tissues, while those genes encoding the more widely expressed subtypes, H1.2, H1.4, and H1x were those bound by more TFs [15]. The proximal promoter of *H1F0* was also targeted by many TFs, in agreement with the plasticity of this subtype, which is known to respond to external stimuli [40] and to compensate H1 decrement upon knockout or knockdown of several subtypes [8, 13].

Correlation analysis of RNA-seq data, from total RNA samples, also divided H1 somatic subtypes into the same two groups. In the first group, transcripts levels of subtypes H1.0, H1.2, H1.4, and H1x were positively correlated. This fact is in agreement with the overlapping detected in the TFs targeting those subtypes. The second group included subtypes H1.1, H1.3, and H1.5, which were negatively correlated with H1.2 and H1.4. These findings suggest possible transcriptional coregulation, which may be associated with the compensatory effects observed in H1 gene knockouts or knockdowns [8][13], and with the normal variability of H1 complement during development and cell differentiation [57]. An argument in favor of the existence of transcriptional co-regulation is that the correlations detected among individual H1 subtypes, at the transcript level, are mostly in agreement with already known biological data, at the protein level. We found that H1.4 has a positive correlation with H1.0 and H1.2 and negative correlation with H1.1, H1.3, and H1.5. It has been described that tissue maturation in mouse is characterized by an increase in H1.0, accompanied by an increase in H1.4 and a decrease in H1.1, and H1.3 [8, 58, 59, 60, 61]. The negative correlation between H1.4 and H1.5 was observed during the differentiation of retinal cells [62]. Although a positive correlation between H1.2 and H1.4 was found at the transcript level, these subtypes have been described to be negatively correlated during maturation of most tissues, except in retina, where both subtypes increased [8, 58, 59, 60, 61, 62]. However, it must be considered that mRNA levels are influenced by post-transcriptional processing, and that translational control may also impact the final protein amount of each subtype.

In addition to the study of the binding of TFs to the proximal promoters, it would be important to identify the enhancers acting on H1 genes and the specific TFs bound to such enhancers. It should be considered that epigenetic regulation, mediated by histone modifications and DNA methylation, may also play a role in the transcriptional regulation of H1 subtypes, especially in diseases like cancer, where epigenetic dysregulation is largely documented. Early studies have shown that the expression of H1.0 increased, after treatment with sodium butyrate, an HDACi, indicating the role of core histone acetylation in *H1F0* transcription [63, 64, 65, 66]. Very recently, a second-generation HDACi, Quisinostat, was able to restore H1.0 expression in cancer cell lines and patient-derived xenografts. The increase in H1.0 expression in tumors was associated with the inhibition of cancer cell self-renewal, and with the anti-tumor properties of the drug [67, 68]. Our results further confirmed the effect of HDAC inhibitors in the expression of H1.0. In addition, we found that the transcript levels of H1x, the other RI-subtype, also increased after treatment with the HDACi, Trichostatin A, while the expression of the RD-subtypes remained relatively constant.

## Conclusions

In summary, in human cell lines, somatic H1 genes were located in the active chromatin compartment with conserved and variable features in their 3D-organization. Interestingly, the region of the histone cluster in chromosome 6 appeared to have more variability in the region assigned to the A compartment, as well as in the chromatin domain length and composition, than the regions containing H1.0 and H1.x genes. The expression of H1 genes was higher when located in TADs, suggesting that intra-TAD promoter-enhancer interactions contribute to their transcriptional activation. Analysis of transcription factors mapped to proximal promoters suggests that somatic H1 genes are under tight redundant and combinatorial regulation during the cell cycle and in different cell types, which may contribute, at least in part, to the variability of H1 complement and may also reflect the functional differentiation of H1 subtypes. We have also found overlapping in the transcription factors targeting H1 promoters and revealed the existence of positive and negative correlations between the mRNA levels of individual subtypes, suggesting co-regulation at transcriptional level. For somatic H1 genes, transcriptional co-regulation did not appear to be determined by physical proximity, as it involves genes located in different chromosomes. Somatic subtypes co-regulation may explain, to some extent, the variability of H1 complement during development, as well as the compensatory effects observed upon H1 knockout or knockdown.

## Materials and Methods

### 3D-chromatin structure data and analysis

Coordinates of the genomic regions containing somatic H1 genes have been extracted from 22 Hi-C experiments, corresponding to 21 human cell lines: hESCs, IMR90 (GSE35156) (27], IMR90, HUVEC, HeLa, NHEK, KBM7, K562, GM12878, HMEC (GSE63525) [26], A549 (GSE105600), Caki2 (GSE105465), G401 (GSE105235), LNCap (GSE105557), NCI-H460 (GSE105725), PANC1 (GSE105566), RPMI-7951 (GSE106022), SCJR (GSE106015), SK-MEL5 (GSE105491), T47D (GSE105697) [29], MCF10a and MCF7 (GSE66733)[28]. Features of each dataset, including the TAD caller, resolution, and boundary definition and length were summarized in Supplementary Table 3. When necessary, all genome coordinates were converted to human genome assembly hg19, using the Liftover tool (UCSC). To group cell lines, hierarchical clustering based on Euclidean distance has been performed with Cluster 3.0, using the coordinates of the chromatin domains. The limits of chromatin domains were considered conserved when their positions shifted no more than 50 kb [26].

### Chromatin compartment analysis

Coordinates of chromatin compartments for 10 human cell lines, at 500 kb resolution, were obtained from bigwig files available at the ENCODE Project [29]. The cell lines used were: A549 (ENCFF460FMH), Caki2 (ENCFF648KNC), G401(ENCFF820URQ), LNCap (ENCFF713UVF), NCI-H460 (ENCFF807HEX), PANC1 (ENCFF783PDP), RPMI-7951 (ENCFF760OPW), SK-MEL5 (ENCFF364HBJ), SK-N-DZ (ENCFF110KIR), T47D (ENCFF411JKH) Overlapping between genomic compartments in the different cell lines was calculated with the Intervene tool [69].

### Transcription factor binding analysis

Data about transcription factor binding have been obtained from the Gene Transcription Regulation Database (GTRD) [31]. To select TF bound to somatic H1 genes we have followed the criteria used by Gokhman and colleagues [34], which considered a gene bound by a TF when the peak of ChIP-seq was within 1000 bp surrounding the transcription start site (TSS) [34]. For further analysis, the initial protein list was filtered, keeping only those proteins considered as human transcription factors [32].

### Gene ontology enrichment analysis

Biological process gene ontology enrichment was performed using Panther Gene Ontology (GO) enrichment analysis tool (http://geneontology.org/). P-values were calculated using Fisher’s Exact Test. Only GO terms with FDR < 0.001 were considered.

### Transcriptomic data

The analysis of H1 somatic expression was performed using RNA-seq datasets from two different RNA sample types: data from polyadenylated mRNA of 1019 cell lines, belonging to the Cancer Cell Line Encyclopedia (CCLE), available at BioProject PRJNA523380, and data from total RNA of 28 human cell lines, available at ENCODE [29]. The specific 28 human cell lines were K562 (GSE88622), HepG2 (GSE88089), GM12878 (GSE78552), IMR90 (GSE90257), A172 (GSE78657), A375 (GSE78652), Caki-2 (GSE78658), Daoy (GSE78636), G401(GSE78662), GM23248 (GSE78670), GM23338 (GSE78682), H1 (GSM438361), H4 (GSE78681), H7 (GSE78651), HT-29 (GSE78684), HT1080 (GSE78653), Karpas-422 (GSE78642), LHCN-M2 (GSE78644), M059J (GSE78666), MG63 (GSE78685), NCI-H460 (GSE78628), OCI-LY7 (GSE78621), PC-3 (GSE78648), RPMI-7951 (GSE78643), SJCRH30 (GSE78655), SJSA1 (GSE78676), SK-MEL-5 (GSE78664) and SK-N-DZ (GSE78627). The GEO accession number for each dataset is indicated in parenthesis. For normalization, the values in transcript per million reads (TPMs) for each subtype were expressed as a percentage of the total TPMs encoding H1 somatic subtypes, in each cell line.

### Correlation analysis between the mRNA levels of H1 somatic subtypes

All the RNA-seq data of the 28 human cell lines, previously described, were used for the correlation analysis. Hierarchical clustering based on the Spearman rank correlation was performed with Cluster 3.0 and visualized with JavaTreeview 1.1.6. Spearman’s correlation coefficients for pairwise subtypes comparison were also calculated. Correlations with p-value < 0.05 were represented in the coexpression network.

### Analysis of the effect of Trichostatin A (TSA) in the expression of H1 somatic subtypes

Breast cancer T47D cells were grown at 37°C with 5% CO2 in RPMI 1640 medium, supplemented with 10% FBS, 2 mM L-glutamine, 100 U/ml penicillin, and 100 g/ml streptomycin. Cells were treated with 400 nM of Trichostatin A for 18 h. Total RNA was extracted using the High Pure RNA Isolation Kit (Roche). The effect of the drug in the expression of H1 somatic subtypes was evaluated by RT-qPCR carried out as previously described [13].

### Statistics

Mann-Whitney U-test analysis, Spearman’s correlation coefficient and Pearson’s correlation coefficient calculations, and statistical significance were carried out using the calculators available at https://www.socscistatistics.com/.

## Declarations

### Availability of data and materials

All the datasets analyzed in this study are available in public repositories: ENCODE, GEO datasets and Bioproject. The accession numbers of all datasets are listed in the methods section.

### Authors’ contributions

IP, analyzed and interpreted Hi-C data, and was a major contributor in writing the manuscript. MA, analyzed and interpreted transcriptomics data. AJ, supplied the experimental data of the effect of HDACi and contributed in writing and revising the manuscript. AR, conceived the study, analyzed the results and wrote most the manuscript. All authors have read and approved the final manuscript.

## Acknowledgements

We thank Andrea Izquierdo-Bouldstridge for her technical help.

## Supplementary files

**Additional file 1. File format: xls**

**Contents: Supplementary Tables 1-8.**

Supplementary Table 1. Coordinates of the A chromatin compartments containing somatic H1 genes in human cell lines.

Supplementary Table 2. Data of the A compartments of chromosomes 3, 6, and 22 in human cell lines.

Supplementary Table 3. Features of the Hi-C datasets.

Supplementary Table 4. 3D-organization of the H1FX gene in chromosome 3.

Supplementary Table 5. 3D-organization of the H1F0 gene in chromosome 22.

Supplementary Table 6. 3D-organization of the somatic H1 genes in chromosome 6.

Supplementary Table 7. Transcription factors mapped to somatic H1 gene promoters.

Supplementary Table 8. Pairwise Spearman’s rank correlation coefficients between the transcript levels of H1 somatic subtypes.

**Additional file2. File format: pdf**

**Contents: Supplementary Figures 1-5.**

Supplementary Figure 1. Length of the A compartment in chromosomes 3, 6, and 22, in human cell lines.

Supplementary Figure 2. Comparison of the Hi-C data of chromosomes encoding somatic H1 genes, from different datasets.

Supplementary Figure 3. Conserved sequence elements in the proximal promoter of somatic H1 genes.

Supplementary Figure 4. Abundance of histone post-translational modifications in somatic H1 gene promoters.

Supplementary Figure 5. Effect of Trichostatin A (TSA) in the transcript levels of H1 somatic subtypes.

