## Additional file 2 for "Towards understanding the regulation of histone H1 somatic subtypes with OMICs"

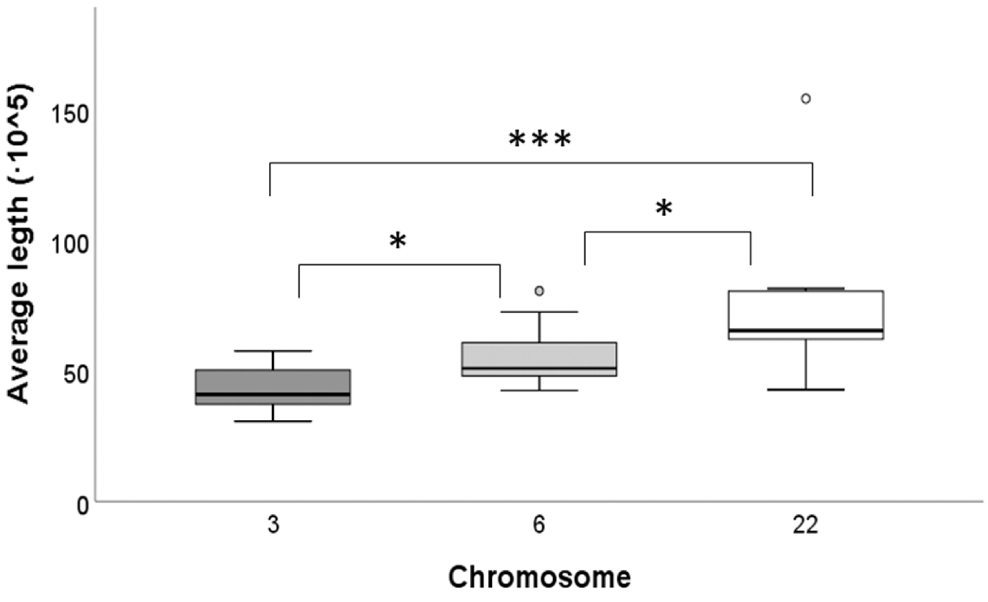


**Supplementary Figure 1. Length of the A compartment in chromosomes 3, 6, and 22 in human cell lines.** Asterisks correspond to the p-values obtained after pairwise comparisons of the length of the A compartments with Mann-Whitney non-parametric test: *, p-value < 0.05; ***, p-value < 0.001.

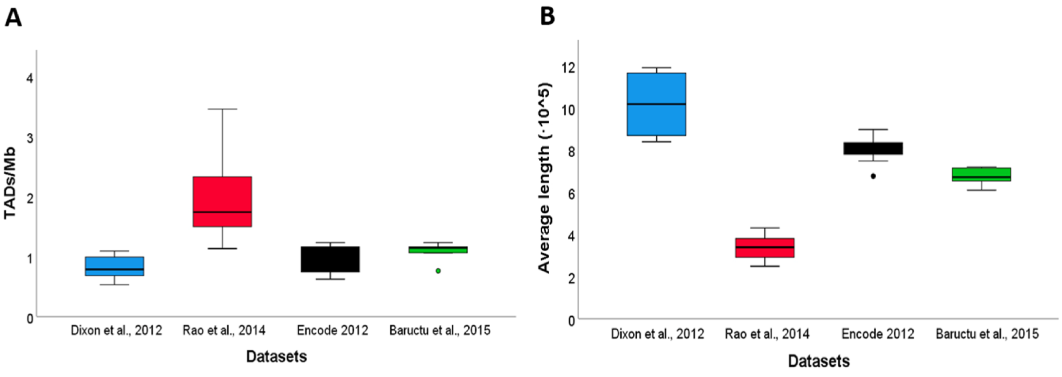


C

p-value of the pairwise comparison of the TADs/Mb between datasets

| Rao et al, 2014 | ENCODE Project 2012 | Baructu et al. 2015 | |
|---|---|---|---|
| 0.0002 | 0.05614 | 0.0455 | Dixon et al., 2012 |
| | 0.00001 | 0.00056 | Rao et al, 2014 |
| | | 0.35238 | ENCODE Project 2012 |

D

p-value of the pairwise comparison of the TADs length between datasets

| Rao et al, 2014 | ENCODE Project 2012 | Baructu et al. 2015 | |
|---|---|---|---|
| 0.0002 | 0.00046 | 0.00512 | Dixon et al., 2012 |
| | 0.00001 | 0.0002 | Rao et al, 2014 |
| | | 0.00512 | ENCODE Project 2012 |

**Supplementary Figure 2. Comparison of the Hi-C data of chromosomes encoding somatic H1 genes from different datasets.** A, Number of TADs per megabase. B, Average length of the TADs. p-values of the pairwise comparisons of the TADs/Mb (C) and TADs length (D) between datasets, with Mann-Whitney non-parametric test. In red p-values < 0.05.

| | TG-Box | H1-Box | |
|---|---|---|---|
| *HIST1H1B* | -531 GTGCC**TGTGTTA**CTTG | -177 CTAAC**AAACACA**ATTT | (354) |
| *HIST1H1C* | -517 GGACC**TGTGTTA**CTTC | -162 GGAAC**AAACACA**ACTT | (355) |
| *HIST1H1D* | -528 GAGCC**TGTGCTA**TTGT | -174 GCAAC**AAACACA**GCAG | (354) |
| *HIST1H1E* | -497 GAGCC**TGTGTTA**CTTC | -143 GTAAC**AAACACA**ACTC | (354) |
| *H1F0* | -344 CCGCG**TGTGTTA**GTTG | +14 GGAAG**AAACACA**GATG | (358) |
| *H1FX* | -451 CGCCC**TGTGTTA**GTTT | -97 CCAAG**AAGCACA**AGTT | (354) |

**Supplementary Figure 3. Conserved sequence elements in the proximal promoter of somatic H1 genes.** In bold, the specific sequence of each element; highlighted in red, the divergent nucleotides within the consensus sequences. In parenthesis, the distance among the two sequence elements. The positions are referred to the transcription start site (TSS).

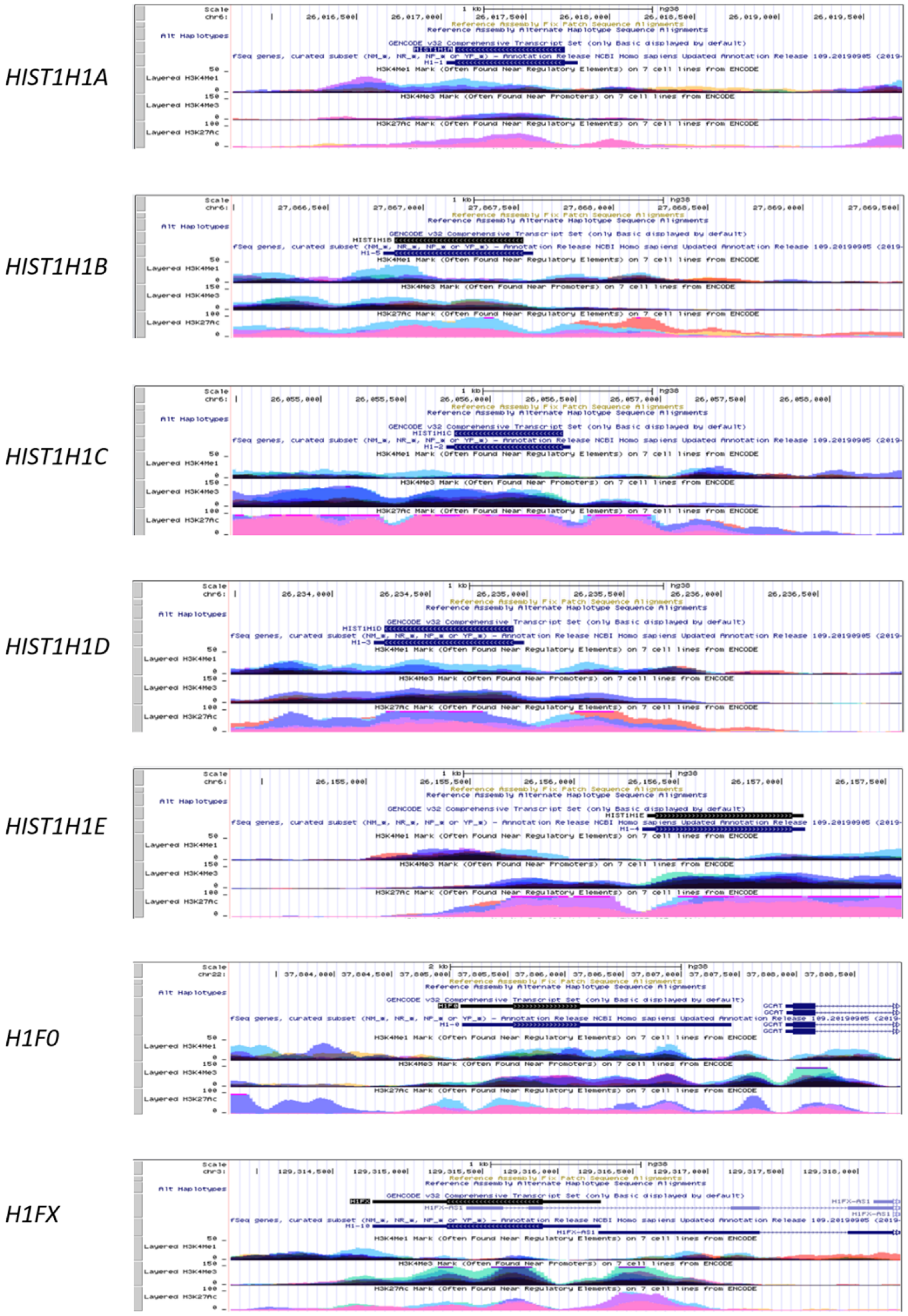


**Supplementary Figure 4. Abundance of histone post-translational modifications in somatic H1 gene promoters.** USCS browser images showing tracks corresponding to H3 modifications, H3K4me1, H3K4me3 and H3K27ac, mapped in 7 human cell lines: GM12878, H1-hESC, HSMM, HUVEC, K562, NHEK and NHLF.

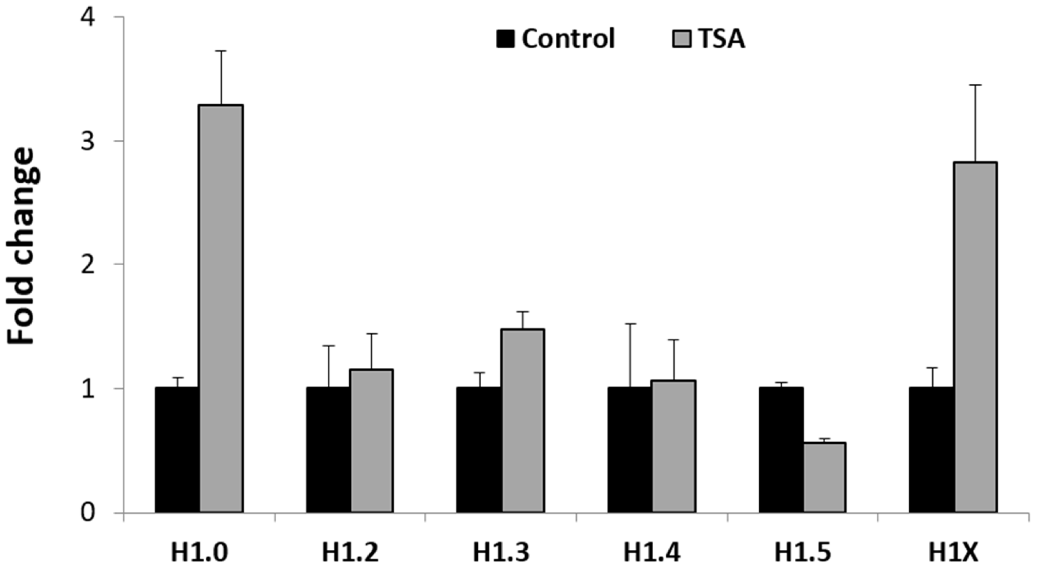


**Supplementary Figure 5. Effect of Trichostatin A (TSA) in the transcript levels of H1 somatic subtypes**. T47D cells were treated with 400nM TSA for 18h. Expression of H1 subtypes was analyzed by RT-qPCR. Fold change correspond to the ratio between the average expression between treated and control samples. Measurements were taken by triplicate, in two independent experiments. Error bars show the standard deviation of the sample.
